# Triple Representation of Language, Working Memory, Social and Emotion Processing in the Cerebellum: Convergent Evidence from Task and Seed-Based Resting-State Fmri Analyses in a Single Large Cohort

**DOI:** 10.1101/254110

**Authors:** Xavier Guell, John D.E. Gabrieli, Jeremy D. Schmahmann

## Abstract

Delineation of functional topography is critical to the evolving understanding of the cerebellum’s role in a wide range of nervous system functions. We used data from the Human Connectome Project (n=787) to analyze cerebellar fMRI task activation (motor, working memory, language, social and emotion processing) and resting-state functional connectivity calculated from cerebral cortical seeds corresponding to the peak Cohen’s d of each task contrast. The combination of exceptional statistical power, activation from both motor and multiple non-motor tasks in the same participants, and convergent resting-state networks in the same participants revealed novel aspects of the functional topography of the human cerebellum. Consistent with prior studies there were two distinct representations of motor activation. Newly revealed were three distinct representations each for working memory, language, social, and emotional task processing that were largely separate for these four cognitive and affective domains. In most cases, the task-based activations and the corresponding resting-network correlations were congruent in identifying the two motor representations and the three non-motor representations that were unique to working memory, language, social cognition, and emotion. The definitive localization and characterization of distinct triple representations for cognition and emotion task processing in the cerebellum opens up new basic science questions as to why there are triple representations (what different functions are enabled by the different representations?) and new clinical questions (what are the differing consequences of lesions to the different representations?).

**HIGHLIGHTS:**

- We analyzed motor and multiple nonmotor task fMRI activations in the cerebellum.
- Resting-state seeds were placed at each task activation peak in the cerebral cortex.
- We describe cerebellar task topography in the largest single cohort studied to date.
- Nonmotor cerebellar task activation revealed a pattern of triple representation.
- Resting-state analysis revealed an overlapping pattern of triple representation.

## 1. INTRODUCTION

Evidence from anatomical, neuroimaging, clinical and behavioral studies indicates that the cerebellum is engaged not only in motor control but also in cognitive and affective functions (Schmahmann, 1991, 1996, 1997; Middleton and Strick, 1994; Schmahmann and Sherman, 1998; Levisohn et al., 2000; Riva and Giorgi, 2000; Ravizza et al., 2006; Schmahmann et al., 2007; Baillieux et al., 2008; Stoodley and Schmahmann, 2009; Thompson and Steinmetz, 2009; Tedesco et al., 2011; Stoodley et al., 2012; Keren-Happuch et al, 2014; Koziol et al., 2014; Hoche et al., 2017). This paradigm shift in appreciation of the clinical neuroscience of the cerebellum has mandated a fundamental reconceptualization of cerebellar organization at the systems level (Schmahmann and Pandya 1997b; Strick et al., 2009; Schmahmann, 2010; Koziol et al., 2014; Mariën et al., 2014; Baumann et al., 2015; Adamaszek et al., 2017).

In the present study, we explored the functional topography of the cerebellum for motor and cognitive functions. This understanding is critical to the Dysmetria of Thought theory and its embedded notion of the Universal Cerebellar Transform. The Dysmetria of Thought theory (Schmahmann, 1991, 1996; 2010; Schmahmann and Sherman, 1998) holds that the cerebellum modulates behavior, maintaining it around a homeostatic baseline appropriate to context. In the same way that cerebellum regulates the rate, force, rhythm and accuracy of movements, so does it regulate the speed, capacity, consistency and appropriateness of mental or cognitive processes. Dysmetria of movement is matched by dysmetria of thought, an unpredictability and illogic to social and societal interaction. The overshoot and inability in the motor system to check parameters of movement are equated, in the cognitive realm, with a mismatch between reality and perceived reality, and erratic attempts to correct the errors of thought or behavior. The theory of the Universal Cerebellar Transform (UCT; Schmahmann, 2000, 2001, 2004) claims that there is a computation unique to the cerebellum because of the essential uniformity of the cerebellar cortical cytoarchitecture (Voogd and Glickstein, 1998; Ito, 1993), and this UCT is applied to all streams of information to which cerebellum has access (Schmahmann, 2000, 2001, 2004; Guell et al., 2017). A corollary of the UCT is the notion of universal cerebellar impairment (UCI), i.e., following cerebellar injury, dysfunction manifests as dysmetria: Dysmetria of movement is the cerebellar motor syndrome; dysmetria of thought and emotion is the cerebellar cognitive affective syndrome (Schmahmann and Sherman, 1998; Levisohn et al., 2000), the third cornerstone of clinical ataxiology (Manto and Mariën, 2015). The Dysmetria of Thought theory is predicated on the existence of two contrasting but complementary anatomic realities: cytoarchitectonic uniformity (the basis of the UCT theory), and highly arranged connectional topography linking distinct cerebellar regions with distinct sensorimotor, association and paralimbic areas of the cerebral hemispheres (Schmahmann and Pandya, 1997a, b; Dum and Strick, 2003; Schmahmann and Pandya, 2008).

The existence and understanding of cerebellar functional topography is thus critical to these contrasting, complementary realities – heterogeneous cerebellar and extracerebellar connectivity, and homogeneous cerebellar cortical cytoarchitecture. Deeper understanding of the presence and arrangement of motor and nonmotor cerebellar functional subregions, the goal of this study, is critical to the evolving understanding of the role of the cerebellum and cerebro-cerebellar interactions in health and disease.

Two motor representations have been recognized in the cerebellum since the work of Snider and colleagues (Snider and Eldred, 1952; see also Dow, 1939; Combs, 1954), one representation in the anterior lobe (lobules IV and V, extending into the rostral aspect of posterior lobe lobule VI) and the other in lobule VIII (**Fig. 1A**). Woolsey, 1952 regarded these as primary and secondary motor representations, along the lines of the dual representation of motor systems in the cerebral hemispheres. This finding has been replicated multiple times: through viral tract tracer studies in monkey in which M1 cerebral cortex injections label cerebellar lobules IV/V/VI and also lobules VIIB/VIII (Kelly and Strick, 2003, **Fig. 1B**), in structure-function correlation studies in patients with stroke (Schmahmann et al., 2009; Stoodley et al., 2016), in PET and task based MRI studies in healthy subjects (Rijntjes et al., 1999; Bushara et al., 2001; Grodd et al., 2001; Takanashi et al., 2003; Thickbroom et al., 2003; Stoodley and Schmahmann, 2009; Buckner et al., 2011; Stoodley et al., 2012; Keren-Happuch et al., 2014), and with resting state functional connectivity MRI (Habas et al., 2009; Krienen and Buckner, 2009; O’Reilly et al., 2010; Buckner et al., 2011). Review of earlier physiological studies in cat (Oscarsson, 1965; see Schmahmann, 2007) demonstrating spinal cord input only to these anterior lobe and lobule VIII regions are consistent with these areas being regarded as the motor cerebellum (Schmahmann, 2004, 2010; Schmahmann et al., 2009; Stoodley et al., 2016).

**Fig.1.**
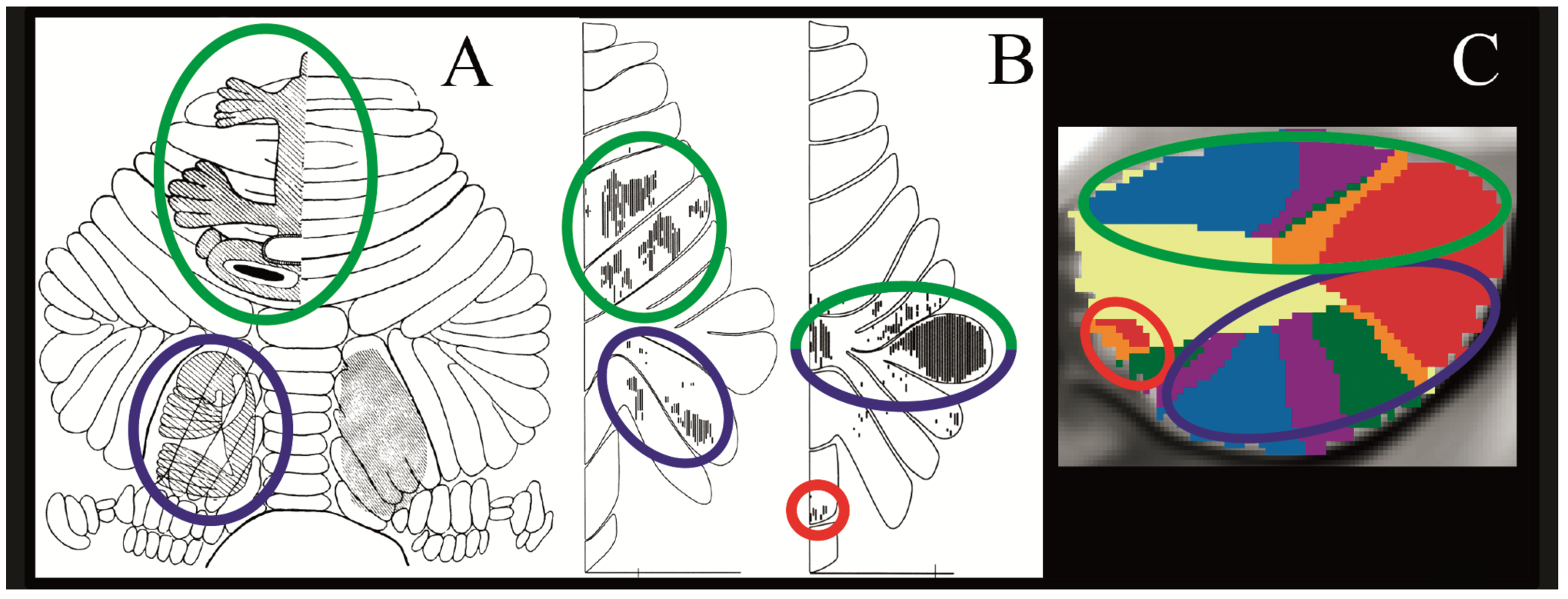
Convergence of findings from multiple studies of cerebellar topography suggesting an overarching organizing principle based on two motor and three nonmotor representations. Green circles indicate first motor (lobules IV/V/VI) and nonmotor (VI/Crus I) representation; blue circles indicate second motor (lobule VIII) and nonmotor (Crus II/VIIB) representation; red circles indicate third nonmotor representation (lobules IX/X). Note that areas of first and second nonmotor representation are contiguous. **A:** Classical electrical stimulation studies showed double representation of sensorimotor activation in the cerebellum (first = lobules IV/V/VI and second = lobule VIII) (Snider and Eldred, 1952; permission pending). **B:** Tract tracing studies demonstrated labeling of the cerebellum in two different locations after injecting viral tracers in motor and nonmotor cerebral cortical areas (viral tracer in M1 labeled cerebellar lobules IV/V/VI and VIIB/VIII, left image; viral tracer in prefrontal cortex area 46 labeled cerebellar lobules Crus I / Crus II and IX, right image) (Kelly and Strick, 2003; permission pending). **C:** Resting-state functional connectivity studies suggest that each resting-state network is represented three times in the cerebellum (approximately lobules IV/V/VI/Crus I, lobules Crus II/VIIB/VIII and lobules IX/X) with the possible exception of the somatomotor network (represented only twice) (image from Buckner et al., 2011, where each color represents one of the seven resting-state networks defined in Yeo et al., 2011).

Knowledge that the cerebellum is engaged in cognition and emotion, and that the anatomical locations of nonmotor cerebellar circuits are different than those for motor circuits emerged first from anatomical tract tracing investigations (Schmahmann and Pandya, 1989, 1991, 1993, 1995, 1997a, b, 2008; Schmahmann, 1996; Middleton and Strick, 1998; Kelly and Strick, 2003; Strick et al., 2009) supported by clinical observations (Schmahmann and Sherman, 1998; Levisohn et al., 2000; Schoch et al., 2006; Schmahmann et al., 2009; Tedesco et al., 2011). Task-based fMRI studies demonstrated that a wide range of cognitive functions activate cerebellum, and a meta-analysis of these studies (Stoodley and Schmahmann, 2009) showed that the cerebellar activations are topographically arranged, an observation supported by a single case of within-individual topography (Stoodley et al., 2010), a prospective study of nine healthy participants (Stoodley et al., 2012), and a second meta-analysis (Keren-Happuch et al., 2014).

Resting-state functional connectivity studies provided additional support for the highly arranged localization within cerebellum of intrinsic connectivity networks subserving different cognitive domains. These studies observed the primary motor representation in the anterior lobe and adjacent lobule VI and the secondary representation in lobule VIII. They also revealed that most of the human cerebellum is not related to cerebral areas involved with sensorimotor processing, but rather is functionally coupled with cerebral association and paralimbic areas. Further, they suggested that there is a triple representation of resting-state cognitive networks in the cerebellum. These three representations localize approximately to (i) lobules VI/Crus I, (ii) lobules Crus II/VIIB/VIII and (iii) lobules IX/X (Buckner et al., 2011; see also Habas et al., 2009; Krienen and Buckner, 2009; O’Reilly et al., 2010) (**Fig. 1C**). Viral tracer studies (Kelly and Strick, 2003) show that area 46 is linked with two of these three areas - lobules Crus II and lobule IX (**Fig. 1B**).A strength of resting-state analyses is that they reflect task-independent correlations among brain regions, but a limitation of such analyses is that they cannot associate specific networks or representations with particular cognitive or emotional functions. Characterizing brain-behavior relations in functional neuroimaging requires tasks that operationalize particular mental operations.

In the present study, we set out to discover the non-motor representational topography of the cerebellum. We aimed for a relatively comprehensive characterization of non-motor domains, examining task-based activations in working memory, language, social cognition, and emotion in the largest single cohort of participants studied to date. We accomplished this by taking advantage of the newly available and unparalleled power in the dataset of the Human Connectome Project with data from 787 participants in the present analysis (HCP; Van Essen et al., 2013). Further, this is the first study to combine the analysis of cerebellar task activation and resting-state functional connectivity in the same group of participants. The advantages of this approach are that (i) resting-state functional connectivity reveals the brain’s intrinsic organization independent of task conditions, (ii) task-activation analyses facilitate the interpretation of the functional significance of topographical patterns and (iii) the combination of the two provides convergent validation of functional topographic maps.

### 2. MATERIALS AND METHODS

#### 2.1 Human Connectome Project data

fMRI data were provided by the Human Connectome Project (HCP), WU-Minn Consortium (Van Essen et al., 2013). EPI data acquired by the WU-Minn HCP used multi-band pulse sequences (Moeller et al., 2010; Feinberg et al., 2010; Setsompop et al., 2012; Xu et al., 2012). HCP structural scans are defaced using the algorithm by Milchenko and Marcus, 2013. HCP MRI data pre-processing pipelines are primarily built using tools from FSL and FreeSurfer (Glasser et al., 2013; Jenkinson et al., 2012; Fischl, 2012; Jenkinson et al., 2002). HCP structural preprocessing includes cortical myelin maps generated by the methods introduced in Glasser and Van Essen, 2011. HCP task-fMRI analyses uses FMRIB’s Expert Analysis Tool (Jenkinson et al., 2012; Woolrich et al., 2001). All fMRI data used in the present study included 2mm spatial smoothing and areal-feature aligned data alignment (“MSMAll”, Robinson et al., 2014).

#### 2.2 Participants

We analyzed data from 787 participants who completed all tasks and resting-state sessions, including 82 couples of monozygotic twins (as determined by genetic testing in the data provided by HCP). 431 were female and 356 were male. Age ranges were distributed as follows: 22-25 (n=172), 26-30 (n=337), 31-35 (n=272), >35 (n=6). HCP exclusion criteria included diabetes or high-blood pressure (for neuroimaging data quality purposes), neurodevelopmental, neuropsychiatric or neurologic disorders, and genetic disorders. Of note, 42 additional subjects are included in the functional connectivity group analysis provided by HCP who were not included in our task activation analysis of 787 participants.

#### 2.3 Tasks and resting-state sessions

HCP provided data from seven tasks (“motor”, “working memory”, “gambling”, “language”, “social”, “emotion” and “relational”), including level 2 cope files for 86 task contrasts (described in Barch et al., 2013 and in Glasser et al., 2016 supplemental materials). We analyzed the following task contrasts: *Movement* (tap left fingers, or tap right fingers, or squeeze right toes, or squeeze left toes, or move tongue) minus *Average* (average of the other four movements), assessing motor function (adapted from Buckner et al., 2011); *Two back* (subject responds if current stimulus matches the item two back) minus *Zero back* (subject responds if current stimulus matches target cue presented at start of block), assessing working memory; *Punish* (money loss blocks) minus *Reward* (money win blocks) and *Reward* minus *Punish*, assessing incentive processing (adapted from Delgado et al., 2000); *Story* (listen to stories) minus *Math* (answer arithmetic questions), assessing language processing (adapted from Binder et al., 2011); *TOM* (view socially interacting geometric objects) minus *Random* (view randomly moving geometric objects), assessing social cognition (adapted from Castelli et al., 2000 and Wheatley et al., 2007); *Relational* (compare featural dimensions distinguishing two pairs of objects) minus *Match* (match objects based on verbal category), assessing relational processing (adapted from Smith et al., 2007); *Faces* (decide which of two angry/fearful faces on the bottom of the screen match the face at the top of the screen) minus *Shapes* (same task performed with shapes instead of faces), assessing emotion processing (adapted from Hariri et al., 2002). Resting-state fMRI data consisted of four 15-minutes scans per subject.

#### 2.4 Analysis of HCP data

We analyzed and visualized data using the Connectome Workbench visualization software and Workbench Command (Marcus et al., 2011). We transformed individual level 2 cope files provided by HCP into Cohen’s d group maps by using ‐cifti-merge followed by ‐cifti-reduce mean, ‐cifti-reduce stdev and ‐cifti-math (cope mean/cope SD). In contrast to level 3 z maps provided by HCP, Cohen’s d maps make it possible to observe the effect size of each task contrast rather than the significance of the BOLD signal change. A sample of 787 subjects ensures that a d value higher than 0.5 (medium effect size, Cohen, 1988) will be statistically significant even after correction for multiple comparisons (given that d=z/sqrt(n), d>0.5 is equivalent to z>14.03 for n=787; analysis of 17,853 cerebellar voxels would require p<0.000028 after Bonferroni correction, and p<0.000028 is equivalent to z>4.026). Accordingly, the analysis did not include any cluster-extent based thresholding as a method of correction for multiple comparisons. This notwithstanding, a cluster size threshold of 100mm3 was used in Table 1 and Fig. 2 - this was done in order to omit very small clusters that would make a comprehensive description of the results too extensive (listing all clusters in Table 1, and labeling all clusters in Fig. 2). Very small clusters were considered to be non-informative for the purpose of a comprehensive characterization of cerebellar functional topography in Table 1 and Fig. 2. In contrast, this cluster size threshold was removed when investigating the possibility of a triple representation of nonmotor function (Fig. 4) – in this portion of the manuscript, inclusion of very small clusters proved to be informative (in particular, in the case of third representation of social processing task activation; see Section 3.3). Of note, because cluster size thresholding was not done for the purposes of statistical significance calculations, removing cluster size threshold in Fig. 4 does not constitute a violation of the methods adopted in our analysis, but rather a full visualization of our results.

**Fig. 2.**
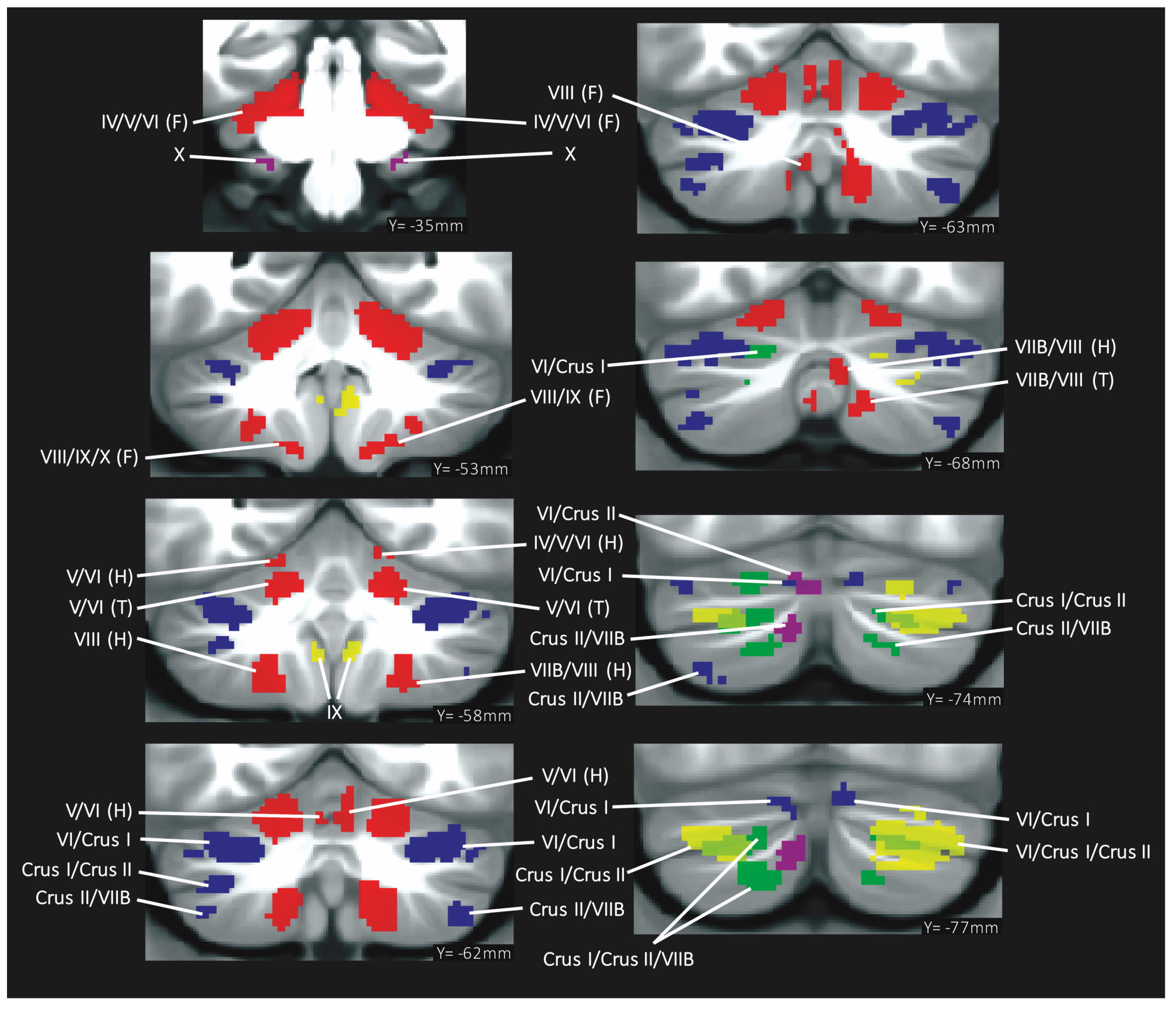
Summary of cerebellar activation for motor (red), working memory (blue), language (yellow), social processing (green) and emotion processing (magenta); coronal plane. (H) = hand, (F) = foot, (T) = tongue. Activation maps are thresholded at a voxel-level threshold of d>0.5. Only clusters >100 mm^3 are shown. Left is shown on the left.

**Table 1.**
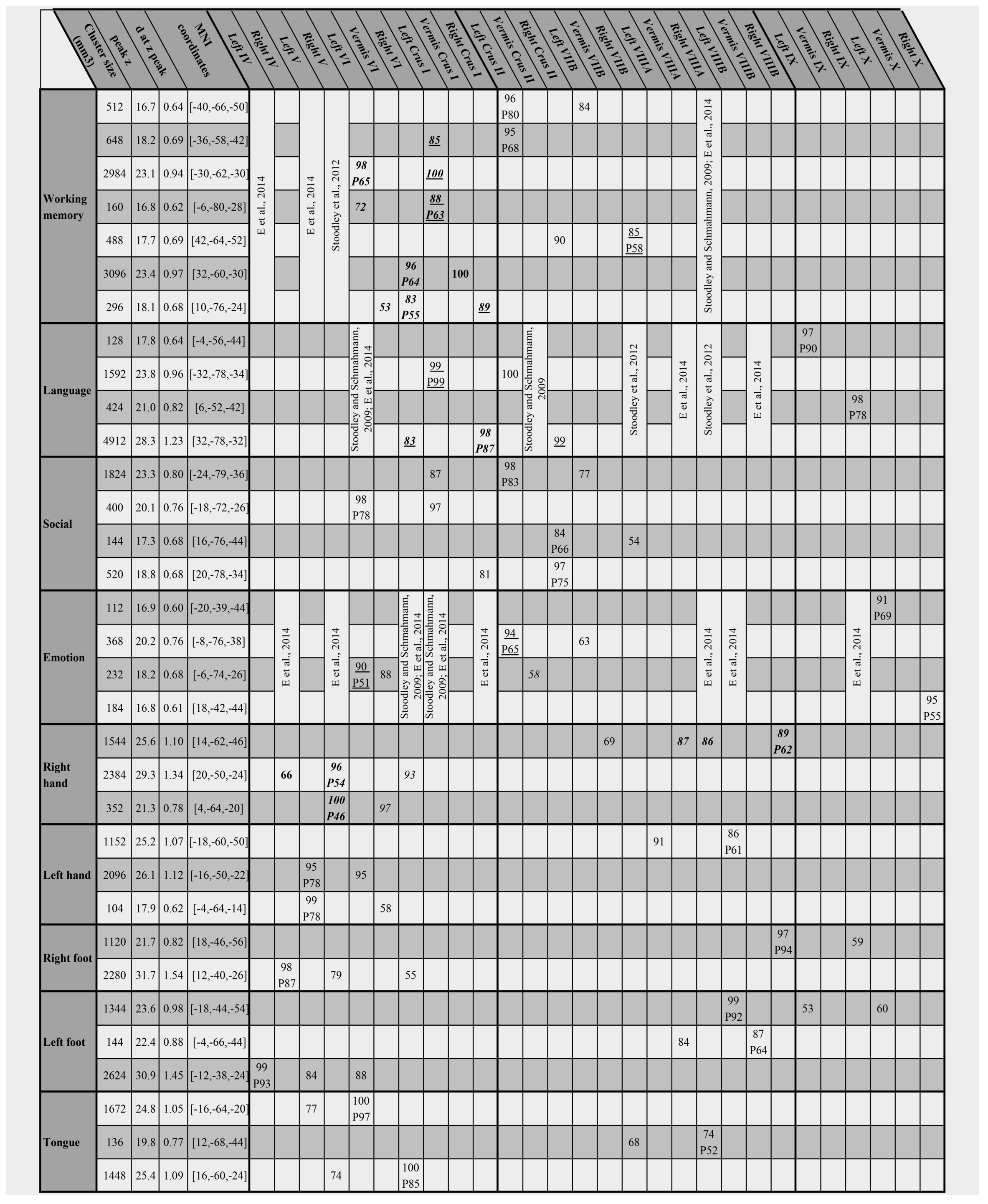
Detailed description of clusters of activation. Key: MNI coordinates = x, y, z coordinates of cluster z peak. Presence of a number written under lobule locations indicates cluster location according to Diedrichsen et al’s FNIRT MNI maximum probability map (Diedrichsen et al., 2009), and value of the number indicates maximum % probability reached at that lobule in the FNIRT MNI probability map. P = z peak location followed by % probability of peak location are also according to the FNIRT MNI maximum probability map (Diedrichsen et al., 2009). Cluster location is written in **bold** if previously described in Stoodley et al., 2012 (group study, n=9), in *italics* if previously described in Stoodley and Schmahmann, 2009 (meta-analysis), underlined if previously described in Keren-Happuch et al., 2014 (meta-analysis), or in regular style if not previously described in any of those three studies.

We identified cerebellar clusters equal to or larger than 100 mm^3 with a d value equal to or higher than 0.5 (medium effect size, Cohen, 1988) using ‐cifti-find-clusters. We calculated the volume of each cluster using ‐cifti-label-import, ‐cifti-all-labels-to-rois and ‐cifti-weighted-stats. Given that HCP uses FNIRT registration to the MNI template, we determined the location of each cerebellar cluster by using Diedrichsen’s FNIRT MNI maximum probability map (Diedrichsen et al., 2009), following the current nomenclature consensus (Schmahmann et al., 2000). While the SUIT probabilistic atlas has shown better overlap between subjects, only 1.75% of the voxels in the cerebellar volume are assigned to a different compartment in FNIRT when compared to SUIT (Diedrichsen et al., 2009). Additionally, we determined the maximum probability reached by each cluster at each lobule by using the FNIRT MNI probability map (Diedrichsen et al., 2009). These maps were downloaded from www.diedrichsenlab.org. Of note, the terms Vermis Crus I and Vermis Crus II are equivalent to the terms lobule VIIAf and lobule VIIAt, respectively (Schmahmann et al., 2000). For ease of reference, figures illustrating anatomical labels and discussion regarding the double/triple representation hypothesis in this article do not distinguish vermal from hemispheric components. In these cases, the terms lobule VI, Crus I, Crus II, lobule VIIB, lobule VIIIA, lobule VIIIB, lobule IX and lobule X refer both to the hemispheric and vermal components of such structures.

We determined the peak z value of each cluster from the level 3 z statistic maps provided by the HCP, as well as the MNI coordinates of the peak z value of each cluster and the corresponding d value as calculated in our analysis. We compared the clusters found after our analysis with the three principal previous reports of motor and nonmotor cerebellar topography (Stoodley and Schmahmann, 2009; Stoodley et al., 2012; Keren-Happuch et al., 2014) and noted whether each cluster location had been previously described in any of these publications.

We visualized resting-state functional connectivity using the group analysis provided by HCP (n=820), and used cerebral cortical seeds corresponding to the peak Cohen’s d of every task contrast. This approach differs from a previous study of cerebellar resting-state networks (Buckner et al., 2011) which applied a winner-takes-all algorithm to determine the strongest functional correlation of each cerebellar voxel to one of the 7 cerebral cortical resting-state networks defined in Yeo et al., 2011. We generated cerebellar maps with a Fisher’s z threshold of 0.309 (given that Fisher’s z = (1/2)[log_e_(1+r) - log_e_(1-r)], Fisher’s z=0.309 is equivalent to r=0.3, and r=0.3 is equivalent to medium effect size [Cohen, 1988]). We overlaid the functional connectivity maps (thresholded at medium effect size, i.e. Fisher’s z>0.309) with task activation maps (thresholded at medium effect size, i.e. Cohen’s d>0.5) to observe whether patterns of task-based functional topography corresponded with patterns of resting-state functional connectivity. Thresholded task activation d maps and thresholded resting-state functional connectivity z maps were also visualized on a cerebellar flat map using the SUIT toolbox for SPM (Diedrichsen, 2006; Diedrichsen et al., 2009; Diedrichsen and Zotow, 2015).

## 3. RESULTS

### 3.1 Clusters of activation of each task contrast

All task contrasts showed cerebellar activation after using a threshold of d>0.5, except for *Relational* minus *Match*, *Punish* minus *Reward* and *Reward* minus *Punish*. Detailed descriptions and illustrations of clusters of activation are included in **Table 1** and **Fig. 2**. **Supplementary Fig. 1 and 2** provide complete coverage of the cerebellum in coronal and sagittal sections.

As in previous studies, we observed motor task activation in lobules IV/V/VI and VIII. In nonmotor tasks, our results replicate previous findings of cognitive task activation in lobule VI and lobule VII (including Crus I, Crus II and lobule VIIB). Analysis also revealed clusters of nonmotor activation in lobules IX and X, an observation which has been previously reported but not always replicated (see Discussion section 4.2.8). Previous studies have revealed encroachment of some nonmotor activations into lobules IV, V and VIII (Stoodley and Schmahmann, 2009; Stoodley et al., 2012; Keren-Happuch et al., 2014). These findings are not reproduced in the large Human Connectome Project cohort data set, providing further support for the selective engagement of these cerebellar lobules in motor rather than nonmotor tasks. Maps of activation thresholded at d>0.5 showed no overlap between motor and non-motor tasks. Further, no overlap was observed within non-motor tasks except between language and social processing.

### 3.2 Location of cerebral cortical seeds for resting-state functional connectivity

Cerebral cortical seeds for resting-state functional connectivity were placed at the peak Cohen’s d of each task contrast, resulting in the following locations (see **Fig. 3**): primary motor cortex (all motor tasks), left angular gyrus (language), right pars triangularis (emotion processing), right superior temporal gyrus (social processing) and right superior parietal cortex (working memory). Note that cerebral cortical regions other than the seeds selected were also engaged in these task contrasts (e.g. left dorsolateral prefrontal cortex was the region with the second highest working memory d value after right superior parietal cortex).

**Fig. 3.**
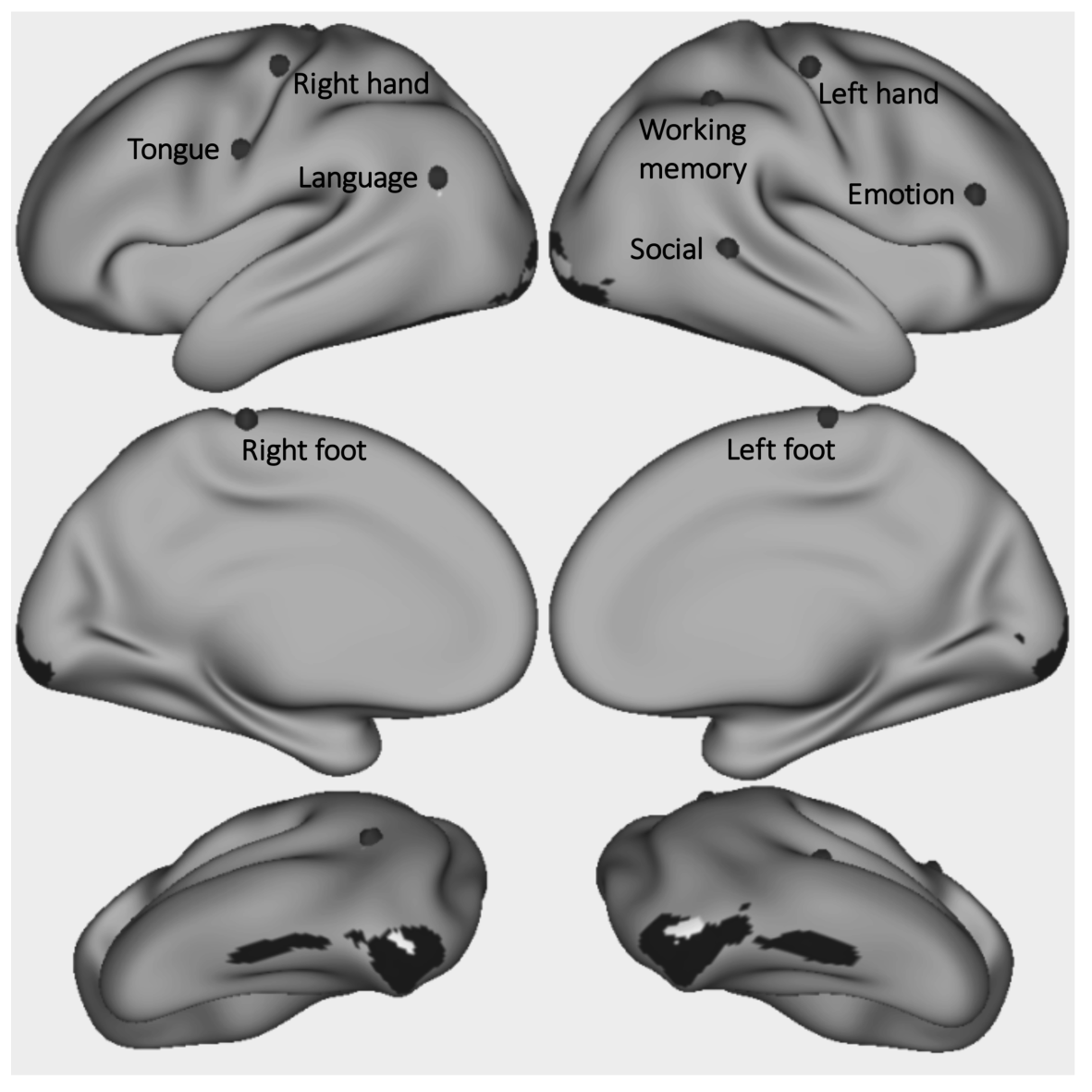
Grey dots mark the seed from which resting-state functional connectivity was calculated for each task contrast. Working memory seed Cohen’s d=1.37. Language d=1.34. Emotion d=1.33. Right hand d=2.75. Left hand d=2.87. Right foot d=2.54. Left foot d=2.62. Tongue d=3.08. Regions shown in black (emotion) and white (social) had a higher task contrast Cohen’s d value than the selected seed location, but resting-state functional connectivity calculated from those seeds did not show overlap with areas of task activation in the cerebellum.

Primary visual cortical areas and left angular gyrus in social processing and primary visual, visual association and inferior temporal gyrus areas in emotion processing had a higher task contrast Cohen’s d values, but seeds from these regions did not show overlap with areas of task activation. These regions are shown in black (emotion) and white (social) in **Fig. 3**. In these cases, emotion and social processing seeds were placed at the next location with the highest Cohen’s d value (right pars triangularis and right superior temporal gyrus, respectively), as indicated in **Fig. 3**.

This is a data-driven approach that did not rely on previous studies of brain function. This notwithstanding, seed locations are consistent with previous reports in the literature of brain function: working memory (e.g. Koenigs et al., 2009), language (e.g. Seghier, 2013), emotion processing (e.g. Dapretto et al., 2006), social processing (e.g. Bigler et al., 2007). Social and emotion processing areas that had a higher task contrast Cohen’s d value but that failed to show overlap with areas of task activation (**Fig. 3**, areas shown in black and white) might correspond to regions involved in primary processing of visual stimuli (primary visual cortex) and face qualities other than emotion (visual association cortex, inferior temporal gyrus; Kravitz et al., 2013).

### 3.3 Double/triple representation hypothesis

We considered the following three hypothetical areas of nonmotor representation as suggested by convergent evidence in the literature (**Fig. 1**): first = lobules VI/Crus I; second = Crus II/VIIB; third = IX/X; in addition to the well-described two areas of motor representation: first = lobules IV/V/VI; second = VIII. When viewing our results within this framework, all motor task contrasts revealed double representation in the cerebellum (**Fig. 4**). Resting-state functional connectivity calculated from cerebral cortical seeds corresponding with Cohen’s d maximum value of each task contrast showed overlap with clusters of task activation, also revealing a pattern of double representation (**Fig. 4**, seeds shown in **Fig. 3**; **Supplementary Fig. 3 - 7** provide full coverage of the cerebellum in sagittal sections and display task activation and resting-state functional connectivity of each motor task contrast, showing overlapping resting-state functional connectivity in all task activation clusters). We did not observe motor task activation in lobules IX or X (third representation) even after removing the cluster size threshold, with the possible exception of foot movement activation extending to lobules IX/X (however, maximum location certainty in lobules IX/X was low [60%], see **Fig. 1**).

**Fig. 4.**
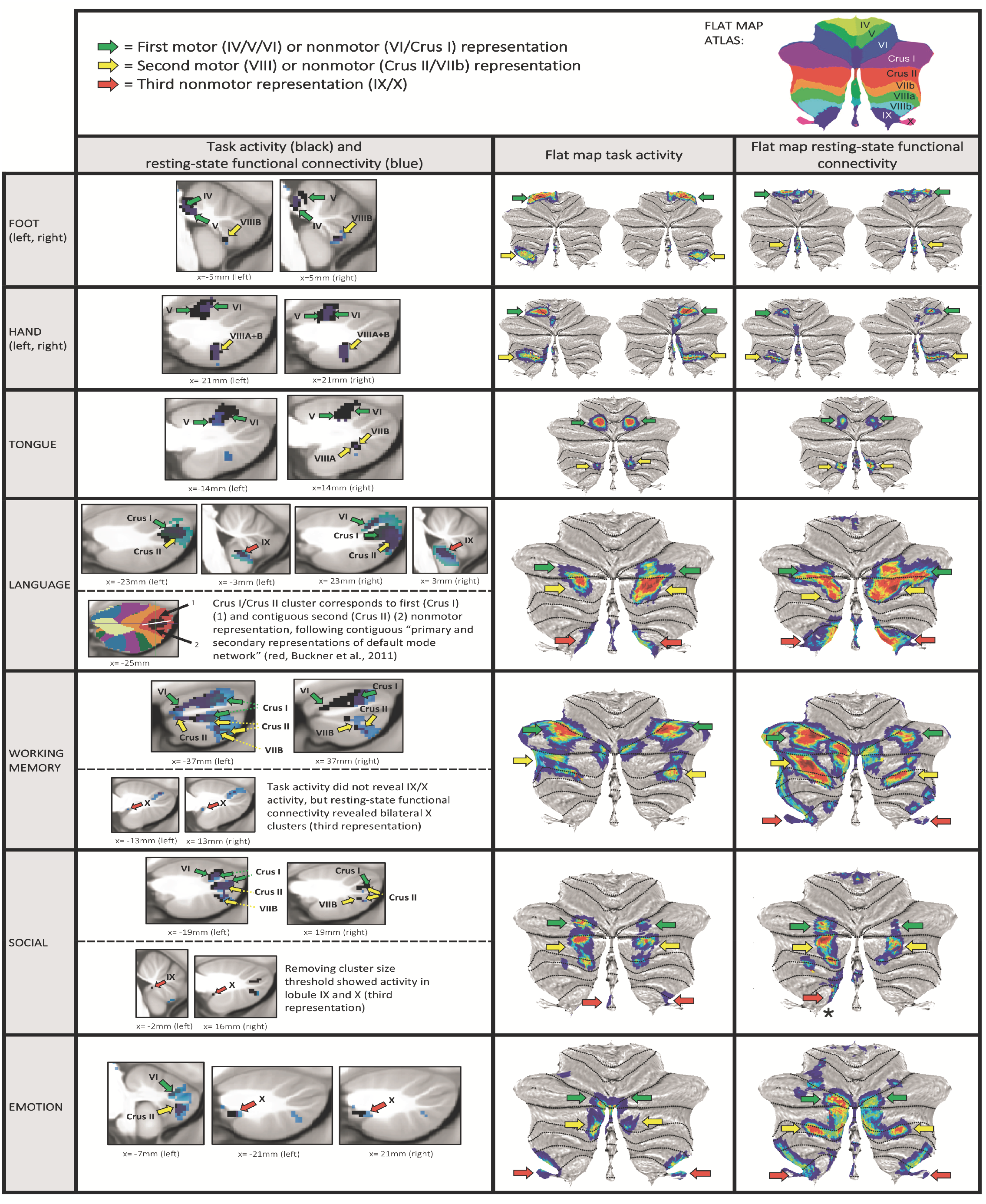
Motor task contrasts revealed double representation (first = lobules IV/V/VI and second = lobule VIII); and nonmotor task contrasts revealed triple representation (first = lobules VI/Crus I; second = lobules Crus II/VIIB; and third = lobules IX/X), except for working memory task activation, which did not reveal a third representation. Resting-state functional connectivity overlapped with clusters of task activation, also revealing a pattern of double motor and triple nonmotor representation. First and second nonmotor representations are sometimes separate (e.g. working memory) and sometimes contiguous (e.g. language processing). Key. First column images: Black = cerebellar task activation (thresholded at d>0.5 [medium effect size] and cluster size>100mm^3). Blue = resting-state functional connectivity calculated from cerebral cortical seed for each task contrast (thresholded at Fisher’s z>0.309, equivalent to r>0.3 [medium effect size]). Second and third column images (flat maps) represent the same resting-state functional connectivity and task activation clusters with no cluster size threshold. Note that cluster size threshold removal does not violate our statistical approach (see section 2.4). Green arrows correspond to first motor or nonmotor representation, yellow arrows correspond to second motor or nonmotor representation, red arrows correspond to third nonmotor representation. Red arrow with an asterisk in social processing resting-state connectivity indicates lobule IX/X engagement that does not overlap with social processing task activation.

All nonmotor task contrasts revealed a pattern of triple representation in the cerebellum (first = lobules VI/Crus I; second = lobules Crus II/VIIB; and third = lobules IX/X) (**Fig. 4**), with the exception of working memory which did not show a third representation (lobule IX/X). Resting-state functional connectivity calculated from cerebral cortical seeds corresponding with Cohen’s d maximum value of each task contrast showed overlap with clusters of task activation, also revealing a pattern of triple representation (**Fig. 4**, seeds shown in **Fig. 3**; **Supplementary Fig. 8 - 11** provide full coverage of the cerebellum in sagittal sections and display task activation and resting-state functional connectivity of each nonmotor task contrast, showing overlapping resting-state functional connectivity in all task activation clusters). The only exception was social processing, which did not reveal a resting-state connectivity cluster overlapping with task activation in the area of third nonmotor representation (lobule IX/X). Clusters extending from Crus I to Crus II were interpreted as contiguous first and second representations, following a previous description of contiguous distribution of primary and secondary representations of the default mode network in Crus I and Crus II (Buckner et al., 2011) (see **Fig. 4,** “language” lower row). Accordingly, first and second nonmotor representations were sometimes separate (e.g. see working memory map in **Fig.4**) and sometimes contiguous (e.g. see language processing map in **Fig. 4**).

In this way, all nonmotor domains revealed triple representation of task activation and seed-based resting-state functional connectivity, with two partial exceptions: (i) social processing revealed triple representation in task activation but not in resting-state connectivity (which did not show an overlapping cluster in the area of third representation [lobule IX/X]); (ii) conversely, working memory revealed triple representation in resting-state functional connectivity but not in task activation (which did not show third representation [lobule IX/X]). This notwithstanding, working memory activation in lobule IX has been previously reported in the literature (Desmond et al., 1997). Further, lowering the thresholding revealed engagement of lobules IX/X in both working memory task activation and social processing resting-state connectivity (see **Supplementary Fig. 12**).

Of note, cluster size threshold (>100mm^3^) had to be removed in order to observe social task activation in lobules IX/X (**Fig. 4,** “social” lower row). This observation agrees with a previous meta-analysis which revealed social processing task activation in lobule IX (Van Overwalle et al., 2014). Note that cluster size threshold removal for social task processing does not violate our statistical approach (see section 2.4).

Notably, while relational and incentive processing did not survive our effect size threshold impositions, these two additional domains also revealed a pattern of triple representation in both task processing and resting-state connectivity when using lower thresholds (see **Supplementary Fig. 13**).

## 4. DISCUSSION

### 4.1 The double motor / triple nonmotor representation hypothesis – relevance and limitations

This is the first study to reveal a pattern of triple and largely distinct representations of cognitive, social, and emotional task processing in the human cerebellum, in addition to replicating the double representation of motor processes in the cerebellum. The triple representation of multiple cognitive, social, and emotional processes was evident in both task activation, which defines the psychological nature of each process, and in resting-state functional connectivity calculated from the peak Cohen’s d of each task activation contrast (**Fig. 4**). Further, the outcomes of the independent cerebellar activation and cortical functional connectivity analyses were largely overlapping – this overlap provides strong convergent evidence for the triple representation of cognitive, social, and emotional functions in the cerebellum, and highlights the relevance of this organization for cortico-cerebellar interactions. Our hypothesis supports and substantially expands Buckner‘s description of triple representation of resting-state networks in the cerebellum (**Fig. 1C**, Buckner et al., 2011). The present findings, from an exceptionally large and high-quality dataset, provide new and fundamental insights into the functional organization of the human cerebellum. Additionally, this study provides a description of cerebellar motor and nonmotor task topography in the largest single cohort of participants studied to date. Activation patterns relative to previous studies are discussed in section 4.2.

We analyzed two aspects of cerebellar physiology – task activation and resting-state functional connectivity. Future studies might investigate whether the same organization applies to other dimensions of anatomical, physiological and pathological cerebellar topography. Previous descriptions in the literature indicate that this might be the case. For example, tract tracing studies in monkeys hint at the possibility of an anatomical correlate of the double motor / triple nonmotor organization: lobules VI/V/VI and VIII receive input from and project to M1, and lobules Crus I/Crus II and IX/X receive input from and project to area 46 (Kelly and Strick, 2003). Similarly, the finding of decreased grey matter in multiple cerebellar locations in autism (Crus I, VIII, and IX), ADHD (IV/Crus I, VIII, and IX) and dyslexia (VI and Crus II) might represent a pathological correlate of the double/triple representation organization (Stoodley, 2014). At a general level, this hypothesis that there is an overarching principle of organization encompassing cerebellar anatomy, physiology and pathology is in accord with current trends in contemporary neuroscience: functional connectivity patterns have been shown to be similar to task activation patterns (Smith et al., 2009), topography of neural degeneration in several diseases seems to follow connectional pathways (Seeley et al., 2009; Zhou et al., 2012; La Joie et al., 2014; Collins et al., 2017), and some resting-state functional networks represent structural connectivity (Greicius et al., 2009; Honey et al., 2009).

Some task contrasts revealed multiple clusters of activation at each area of representation. For example, working memory revealed two clusters in lobules VI/Crus I bilaterally, all four corresponding to first representation according to our hypothesis; and two left and one right Crus II/VIIB clusters, all three corresponding to second representation according to our hypothesis. Multiple clusters of activation in one cerebellar region might appear to conflict with the notion that all activation within that area corresponds to the same representation of a given domain. However, multiple clusters of activation have also been observed within the well-established primary and secondary areas of representation of motor activation in the cerebellum. For instance, our analysis revealed two clusters both corresponding to second representation of left foot movement (lobule VIII), two clusters both corresponding to first representation of left hand movement (V/VI), and two clusters both corresponding to first representation of right hand movement (IV/V/VI).

The HCP imaging protocol has the strength of being exceptionally comprehensive in measuring multiple domains, but also the limitation that each domain is assessed in a brief and broad fashion. For example, language processing is defined as a contrast between story listening versus math performance, and such a contrast involves many specific language processes including semantics, syntax, phonology, and pragmatics. Future studies will be needed to evaluate language and other domains in greater analytic detail. The comprehensive range of domains examined, however, is well suited to discovering broad organizational principles of cerebellar functions that may be domain-independent. Of note, while relational and incentive processing did not survive our effect size threshold impositions, these two additional domains also revealed a pattern of triple representation in both task processing and resting-state connectivity when using lower thresholds (**Supplementary Fig. 13**). The observation of a triple representation of multiple nonmotor domains in both task activation and resting-state connectivity supports the view that the triple nonmotor organization reflects a general property of cognitive and affective processing in the cerebellum, rather than a distribution unique to the particular task contrasts included in the present study.

The biological significance of the hypothesized double/triple representation organization remains to be determined. Our use of the terms “first”, “second” and “third” representation does not necessarily imply an analogy with, for example, the multiple cerebral cortical representations of motor function (Woolsey et al., 1952). This notwithstanding, clinical evidence suggests that the importance of the first and second representations of motor function in the cerebellum is not equal - lesion of lobules IV/V/VI results in more severe motor deficits than do lesions of lobule VIII (Schmahmann et al., 2009; Stoodley et al., 2016). Similarly, the observation of working memory task activation and social processing resting-state connectivity only when lowering our threshold impositions (**Supplementary Fig. 12**) hints at the possibility that engagement of lobules IX/X in nonmotor processes might be more prominent in some cases. Future studies might investigate whether an asymmetry exists between the physiological and pathophysiological contributions of the apparent first, second and third representations of nonmotor functions in the cerebellum.

It did not escape our notice that the organization along a common sagittal axis of multiple representations of task activation of each task contrast (**Fig. 2**, **Supplementary Fig. 1**) is reminiscent of the organization along the parasagittal axis of the well-established molecular compartmentalization in the cerebellum, the zebrin stripes (Brochu et al., 1990; Oberdick et al., 1998; Voogd and Glickstein, 1998). This parasagittal organization also seems to be obeyed by corticonuclear and olivocerebellar projection fibers in the cerebellum (see figure 3A in Voogd and Glickstein, 1998; also Sugihara and Shinoda, 2004). Therefore, it is a possibility that the pattern of multiple representations described here respects the sagittal organization of cerebellar fibers, extending vertically and therefore encompassing lobules IV/V/VI/Crus I (first motor and nonmotor representations), Crus II/VIIB/VIII (second motor and nonmotor representations) and IX/X (third nonmotor representation).

The nature of the contribution of the cerebellum to movement, thought and emotion is still being elucidated (Schmahmann, 1997, 2010; Koziol et al., 2014; Mariën et al., 2014; Baumann et al., 2015; Adamaszek et al., 2017), and the cerebellar structural and functional abnormalities in diseases such as depression, anxiety, bipolar disorder, schizophrenia, dyslexia, autism and ADHD remain areas of active investigation (Phillips et al., 2015). While still at an initial stage of development, the double/triple representation hypothesis might critically influence future studies investigating the anatomy, physiology and neuropsychiatry of the cerebellum and corticosubcortical interactions. One immediate application of the double/triple representation hypothesis is in the interpretation of cerebellar cognitive, social or affective task activation findings in healthy subjects or patient populations. Neuroimaging studies analyzing cerebellar task activation commonly report motor findings in terms of “first motor representation” (lobules IV/V/VI) and “second motor representation” (lobule VIII). Cognitive, social and affective task activation clusters have never been reported, to our knowledge, in terms of three representations (lobule IV/Crus I, Crus II/lobule VIIB, and lobule IX/X) - it is reasonable to consider that this conception could crucially and immediately influence future interpretations of cerebellar task activation findings. We propose that just as the double motor representation has become the common clinical and scientific framework for interpreting motor functions of the cerebellum, now the triple non-motor representation should become the common clinical and scientific framework for interpreting the cognitive, social, and emotional functions of the cerebellum.

### 4.2 Cerebellar functional topography

This study provides a description of cerebellar motor and nonmotor task topography in the largest single cohort of participants studied to date. For this reason, in this section we provide an extensive description and discussion of activation patterns relative to previous studies, with a focus on the three principal previous reports of motor and nonmotor cerebellar topography (two meta-analyses by Stoodley and Schmahmann, 2009 and Keren-Happuch et al., 2014; and a group study by Stoodley et al., 2012, n=9).

#### 4.2.1 Motor versus nonmotor regions

A differentiation of motor versus nonmotor regions in the cerebellum has been supported by anatomical and clinical observations - evidence suggests that while lobules IV, V, VI and VIII are engaged in motor tasks, nonmotor processing occurs in lobules VI and VII. Projections carrying spinal cord input terminate in lobules IV, V and VIII, whereas projections carrying no spinal cord input terminate in lobules VI and VII (as reviewed in Schmahmann, 2007; see also Stoodley and Schmahmann, 2010). Similarly, while motor cortex is linked with lobules IV, V and VI, prefrontal cortex is linked with lobule VII (Kelly and Strick, 2003). Also, motor symptom presentation correlates preferentially with lesions in lobules IV, V and VI (Schoch et al., 2006; Schmahmann et al., 2009; Stoodley et al., 2016). While all three principal previous reports of motor and nonmotor cerebellar topography observed nonmotor activation which sometimes extended to lobules IV, V or VIII (Stoodley and Schmahmann, 2009, working memory activation in lobule VIII; Stoodley et al., 2012, working memory activation in lobule V and language processing activation in lobule VIII; Keren-Happuch et al., 2014, working memory activation in lobule IV, V and VIII, language processing activation in lobule VIII and emotion processing activation in lobules IV, V and VIII), our analysis did not reveal cerebellar nonmotor activation in any of these locations. In this way, our findings strongly support the motor specificity of cerebellar lobules IV, V and VIII.

#### 4.2.2 Independent representations of nonmotor regions

For the most part, there were distinct cerebellar activations for working memory, language, emotion, and social cognition. Although any claim for distinct representations necessarily rests upon a selected threshold, the absence of an activation with over 700 participants is highly suggestive of domain-specific representations of different kinds of cognition and emotion in the cerebellum. The exception was an overlap between activations for language and social tasks, perhaps reflecting similarities between cognitive processes engaged in these particular assessments of language (story listening) and social cognition (viewing socially interacting geometric shapes, possibly involving similar narrative components). Contrasting with previous studies reporting an overlap between cerebellar social processing activation and other executive and semantic functional regions (Van Overwalle et al., 2014), our observation supports a cerebellar domain-specific contribution to social cognition (as argued more recently in the reinterpretation of Van Overwalle et al., 2014 in Van Overwalle et al., 2015).

#### 4.2.3 Motor tasks

Motor task lateralization was evident. Right hand and foot tasks engaged the right cerebellum only, while left hand and foot tasks engaged the left cerebellum only. Tongue movements generated bilateral cerebellar activation.

As in previous reports (Stoodley et al., 2012; Stoodley and Schmahmann, 2009; see also Snider and Eldred, 1952; Rijntes et al., 1999; Bushara et al., 2001; Grodd et al., 2001; Takanashi et al., 2003; Thickbroom et al., 2003), right hand movement generated activation in right lobules IV, V, VI and VIII. Of note, lobule VI motor activation was observed as an extension of lobule V motor activation and did not overlap with areas of cognitive task activation in lobule VI (see **Fig. 2**), further supporting the notion that distinct motor and cognitive functional subregions exist in the cerebellum even within lobules which are engaged in both motor and cognitive processing. All other motor tasks revealed a similar IV/V/VI and VIII distribution.

#### 4.2.4 Working memory task

Cerebellar engagement in working memory tasks is reliably reported across multiple fMRI and PET studies (e.g. Fiez et al., 1996; Desmond et al., 1997; Beneventi et al., 2007; Hautzel et al., 2009; Hayter et al., 2007; Kirschen et al., 2005, 2010; Marvel and Desmond, 2010), and numerous publications report working memory deficits after cerebellar injury (Schmahmann and Sherman, 1998; Silveri et al., 1998; Ravizza et al., 2006; Peterbus et al., 2010).

In agreement with the three principal previous reports of motor and nonmotor cerebellar topography (Stoodley and Schmahmann, 2009; Stoodley et al., 2012; Keren-Happuch et al., 2014), our analysis revealed activation in right and left Crus I. We also found left and right lobule VI activation, a finding that had been previously reported in Stoodley and Schmahmann, 2009 and Stoodley et al., 2012, but that was not replicated in Keren-Happuch et al., 2014. Our results also indicate activation in right lobule VIIB, a finding previously reported in Keren-Happuch et al., 2014. Additionally, our analysis showed cerebellar engagement in right Crus II, left Crus II and lobule VIIB; these areas had not been previously described in any of the three principal previous studies of motor and nonmotor cerebellar topography but are coherent with the well-established cognitive function of these regions (e.g. language, social cognition, spatial processing, executive function; see Stoodley and Schmahmann, 2009; Stoodley et al., 2012; Keren-Happuch et al., 2014).

#### 4.2.5 Language processing task

Cerebellar injury has been shown to result in a constellation of language deficits. The original description of the CCAS (Schmahmann and Sherman, 1998) reported agrammatism, dysprosodia, anomia, verbal fluency and verbal working memory deficits; and later investigations also revealed metalinguistic deficits (Guell et al., 2015) (see Mariën et al., 2014 for a review). Clinical (Scott et al., 2001; Gottwald et al., 2004) and neuroimaging findings (Fiez and Raichle, 1997; Hubrich-Ungureanu et al., 2002; Jansen et al., 2005) support rightlateralization of language function in the cerebellum, and our analysis also revealed wider (cluster size) and stronger (Cohen’s d) language activation in the right cerebellum. Consonant with the three principal previous reports of motor and nonmotor cerebellar topography (Stoodley and Schmahmann, 2009; Stoodley et al., 2012; Keren-Happuch et al., 2014), we found cerebellar engagement in right lobule VI. We also found activation in right Crus I, a finding that had been reported in Stoodley and Schmahmann, 2009 and Stoodley et al., 2012 but that had not been replicated in Keren-Happuch et al., 2014. Analysis also revealed activation in left Crus I and right Crus II, as in Keren-Happuch et al., 2014. Further, we observed right and left lobule IX and left Crus II activation. These areas had not been previously described in any of the three principal previous studies of motor and nonmotor cerebellar topography (Stoodley and Schmahmann, 2009; Stoodley et al., 2012; Keren-Happuch et al., 2014), but nonetheless have a well-established role in nonmotor processes (see section 4.2.8 for a discussion on lobule IX). While Stoodley and Schmahmann, 2009 and Keren-Happuch et al., 2014 had previously reported activation in left lobule VI and Crus II vermis, our analysis did not replicate this finding.

#### 4.2.6 Social processing task

Social processing was not investigated in the three principal previous studies of motor and nonmotor cerebellar topography (Stoodley and Schmahmann, 2009; Stoodley et al., 2012; Keren-Happuch et al., 2014). This notwithstanding, cerebellar social processing task activation has been reported in many previous PET and fMRI studies, as reviewed in a recent meta-analysis by Van Overwalle et al., 2014. Notably, cerebellar injury has been shown to result in lack of empathy as well as difficulty with social cues and interactions (Schmahmann et al., 2007; Garrard et al., 2008; Hoche et al., 2016).

Our analysis revealed bilateral activation in Crus I, Crus II and VIIB as well as in left lobule VI - these observations match those in the meta-analysis by Van Overwalle et al., 2014, which localized the majority of clusters of cerebellar social processing task activation in lobules VI, Crus I and II. Van Overwalle et al., 2014 also identified a cluster in lobule IX, which was also found in our analysis after removing the cluster size threshold (>100mm^3^).

#### 4.2.7 Emotion processing task

Cerebellar engagement in emotion processing has been supported by functional neuroimaging (Lane et al., 1997; Schraa-Tam et al., 2012; Baumann and Mattingley, 2012), TMS (Schutter et al., 2009; Schutter and Van Honk, 2009; Moulton et al., 2010) and clinical studies (Schmahmann and Sherman, 1998; Schmahmann et al, 2007; Turner et al., 2007). Following the meta-analysis of Keren-Happuch et al., 2014, our analysis also revealed activation in left Crus II and left lobule VI, as well as in Crus II vermis as reported in the meta-analysis of Stoodley and Schmahmann, 2009.

We also observed cerebellar engagement in right and left lobule X, an area of activation which had been previously reported in a study using a negative faces vs neutral faces task contrast (Schraa-Tam et al., 2012). However, many other studies failed to identify lobule X emotion task activation (Stoodley and Schmahmann, 2009; Stoodley et al., 2012; Keren-Happuch et al., 2014; Baumann and Mattingley, 2012). The nature of the emotional stimuli may explain this apparent contradiction between studies. The emotional task in our analysis included angry and fearful faces, resembling the stimuli used in Schraa-Tam et al., 2012. In contrast, emotion tasks in other studies not revealing lobule X activation included a diverse range of emotions. Neuroimaging and cerebellar patient studies suggest that the cerebellum is engaged by negative rather than positive emotions (Turner et al., 2007; Baumann and Mattingley, 2012). The use of angry and fearful face expressions might therefore explain our observation of lobule X activation, and suggests that future neuroimaging task studies aiming to engage the limbic cerebellum may benefit from using negative valence emotion stimuli.

Given that the substrate of the cerebellar-vestibular system has long been thought to be lobule X, our finding suggests a relationship between the cerebellar-vestibular system and cerebellar emotion processing. This relationship was proposed by Schmahmann (1991): “*(O)lder cerebellar regions consisting of the flocculonodular lobe, vermis, and associated fastigial and— to a lesser degree—globose nuclei, could perhaps be considered as the equivalent of the limbic cerebellum…concerned with primitive defense mechanisms including the autonomic manifestations of the fight or flight response, as well as with emotion, affect, and sexuality and, possibly, also with affectively important memory*”. In line with this hypothesis, Schmahmann et al., 2007 reported two cases of patients with cerebellar injury including lobule X who presented with a vestibulocerebellar disorder as well as panic episodes - in both cases, panic episodes were precipitated and exaggerated by motion. Additionally, Levinson, 1989 observed 93% of prevalence of at least one cerebellar-vestibular sign in the electronystagmographic examination of patients diagnosed with an anxiety disorder (n=402), including panic disorder, generalized anxiety disorder, post-traumatic stress disorder, social phobia and specific phobia. The fact that emotion processing activates lobule X suggests a potential shared anatomical substrate of emotion processing and vestibular function, and thus potentially explains anxiety in patients with lesions of the vestibulocerebellum (lobule X), as well as cerebellar-vestibular abnormalities in patients with primary anxiety disorders.

As in previous reports (Stoodley et al., 2009; Baumann and Mattingley, 2012; Keren-Happuch et al., 2014), we identified emotion processing task activation at the cerebellar vermis (see **Fig. 4**). This observation supports the vermal “limbic cerebellum” hypothesis (Schmahmann, 1991, 2000, 2004), which draws on the connections of this structure with limbic brain areas (Schmahmann, 1996), reports of modulation of emotion following cerebellar midline manipulation or stimulation in animals and humans (Heath, 1977; Berman et al., 1978; Levisohn et al., 2000), as well as with clinical observations revealing vermal involvement in patients with pronounced affective symptoms (Schmahmann and Sherman, 1998; Levisohn et al., 2000; Gudrunardottir et al., 2016).

#### 4.2.8 Nonmotor cerebellar activation in lobules IX and X

Lobule IX is considered important for visual guidance of movement, and vestibular function is attributed to lobule X (Voogd et al., 1996; Stoodley and Schmahmann, 2010). With the exception of emotion processing activation in lobule IX (Keren-Happuch et al., 2014), nonmotor task activation has not been reported in lobules IX or X in any of the three principal previous studies of motor and nonmotor cerebellar topography (Stoodley and Schmahmann, 2009; Stoodley et al., 2012; Keren-Happuch et al., 2014). In contrast, our analysis revealed lobule IX/X nonmotor activation in multiple nonmotor task activation and resting-state connectivity maps. These observations are harmonious with previous studies reporting nonmotor lobule IX activation (e.g. working memory, Desmond et al., 1997; timing perception, Liu et al., 2008; emotion processing, Schraa-Tam et al., 2012); as well as default-mode network representation in lobule IX (Habas et al., 2009; Krienen and Buckner, 2009; O’Reilly et al., 2010; Buckner et al., 2011) and representation of dorsal attention and frontoparietal networks in lobule X (Buckner et al., 2011).

#### 4.2.9 Task contrasts which did not reveal cerebellar activation

Although previous studies have shown cerebellar activation in reward (Völlm et al., 2007; Nees et al., 2012; Shigemune et al., 2014; Kahn et al., 2015, Katahira et al., 2015) and loss/punishment tasks (Völlm et al., 2007; White et al., 2014), our analysis and Cohen’s d threshold of d>0.5 did not reveal cerebellar activation in the *Punish* minus *Reward* or *Reward* minus *Punish* task contrasts (maximum cerebellar d=0.15 and d=0.26, respectively).

In the reasoning “*relational*” condition, participants had to infer the dimension of difference between two objects (e.g. shape or texture) and then infer whether two other objects differed along the same dimension. In the control (“*match*”) condition, participants saw three objects and had to infer whether the third object matched any of the first two objects in a specified dimension (shape or texture). In this way, the *Relational* minus *Match* task contrast assessed the cognitive process of relational matching, which includes reasoning and working memory processes (Smith et al., 2007). Ample evidence supports a cerebellar role in such functions (e.g. Schmahmann and Sherman, 1998; Stoodley et al., 2009; Keren-Happuch et al., 2014; and also this present study), but maximum cerebellar Cohen’s d value for *Relational* minus *Match* was 0.45 in our analysis. As an attempt to display clusters of activation which are not only statistically significant in the context of a large sample size, we established a Cohen’s d threshold of 0.5 (medium effect size, Cohen, 1988) and therefore did not show functional task topography for the punish/reward and relational task contrasts. Lower thresholding in these tasks is reported in Supplementary Fig. 13.

## 5. CONCLUSION

We show for the first time that there is a triple representation of nonmotor task activation in the cerebellum. A resting-state analysis from seeds placed at each task activation peak in the cerebral cortex revealed an overlapping pattern, providing strong convergent evidence for the double motor / triple nonmotor organization. These findings unmask novel and fundamental questions which might become critical for the understanding of cerebellar physiology and pathophysiology at the systems-level. It is known that cerebellar damage to the first motor representation results in motor deficits more severe than after damage to the second motor representation (Schmahmann et al., 2009; Stoodley et al., 2016) - what are the distinct consequences of injury to the areas of first, second and third nonmotor representation? Do structural and/or functional abnormalities in psychiatric diseases map to any of these areas preferentially? What is the contribution of each representation to nonmotor processing in the cerebellum? - can a task contrast analysis demonstrate preferential engagement of a particular nonmotor representation (first, second or third) for a particular task characteristic, perhaps consistently across cognitive domains? What is the relationship between the two motor and three nonmotor representations? - do any asymmetries (physiological or pathophysiological) between the first and second motor representations extrapolate to their adjacent first nonmotor and second/third nonmotor representations? In many ways the discovery of three task processing representations in each of multiple non-cognitive domains opens up more questions than it answers, but this discovery also establishes a fundamental organizational principle of the human cerebellum that ought to be a guiding framework for future clinical and basic science research.

## ACKNOWLEDGMENTS

We thank Susan Whitfield-Gabrieli, Satrajit S. Ghosh and Oscar Vilarroya for discussion.

